# HIV-1 controllers possess a unique CD8+ T-cell activation phenotype and loss of control is associated with increased expression of exhaustion markers

**DOI:** 10.1101/2024.04.09.588737

**Authors:** Amber D Jones, Zachary Capriotti, Erin Santos, Angel Lin, Rachel Van Duyne, Stephen Smith, Zachary Klase

**Affiliations:** Department of Biological Sciences, University of the Sciences, Philadelphia, PA 19104, USA; Department of Pharmacology and Physiology, Drexel University College of Medicine, Philadelphia, PA 19102, USA; Molecular and Cell Biology and Genetics Graduate Program, Drexel University College of Medicine, Philadelphia, PA, USA; The Smith Center for Infectious Diseases and Urban Health, West Orange NJ 07018, USA; Center for Neuroimmunology and CNS Therapeutics, Institute for Molecular Medicine and Infectious Diseases, Drexel University College of Medicine, Philadelphia, PA 19102; Sidney Kimmel Cancer Center, Thomas Jefferson University, Philadelphia, PA, 19102

## Abstract

HIV-1 controllers are a rare population of individuals that exhibit spontaneous control of HIV-1 infection without antiretroviral therapy. These controllers can be categorized based on the level and mechanism of control. Understanding the mechanisms by which controllers maintain and eventually lose this ability would be highly valuable in HIV-1 cure or vaccine research. We explored whether CD8+ T cell exhaustion plays a role in the maintenance and loss of control by examining immune characteristics of HIV-1 controllers and controllers who lost control within the duration of the study. Previous work revealed the ability of CD8+ T-cells isolated from HIV-1 controllers to suppress HIV-1 replication in matched CD4+ T-cells *ex vivo*. Using flow cytometry, we analyzed exhaustion marker expression on CD8+ T-cells from these controllers and determined that they maintain a unique exhaustion profile as compared to HIV-negative individuals and standard progressors. The low level of T-cell exhaustion seen in controllers was reversed when these individuals lost control and showed increased viral loads. Immune checkpoint blockade targeting exhaustion markers was able to restore *ex vivo* control by CD8+ T-cells from former controllers. These results suggest that CD8+ T cell exhaustion compromises the ability to control viral replication in HIV-1 controllers.

**AUTHOR SUMMARY:** Despite the use of antiretroviral therapy, HIV continues to be a major public health issue that affects the lives of millions of people. Some infected people can control viral infection without therapy. The mechanism by which some people can control HIV infection at low, but detectable levels is unknown. In this study we examined the state of cytotoxic CD8+ T-cells in a group of HIV controllers and found that controllers maintain low levels of markers for a chronic state of activation called exhaustion. Loss of control correlated with increase in exhausted T-cells and for individuals who had recently lost control of infection we could restore protection in the cell culture dish by using immune checkpoint blockade drugs that affect exhaustion.

## INTRODUCTION

CD4+ T cells are among the main cell types targeted by HIV-1 *in vivo* (1). In the standard progression of HIV-1 without therapy, normal progressors experience a decline in immune function (2). Within the first 6-8 weeks of HIV-1 infection, the host experiences a significant decrease in CD4+ T cells, a concomitant increase in viral load, and a corresponding increase in immune activation (3–5). To control the infection, a CD8+ T cell-mediated adaptive immune response is mounted (6, 7). Despite these efforts, the virus is not eliminated and due to sustained antigen production, CD8+ T cells undergo a state of T cell exhaustion characterized by reduced functionality that can be detected via increased expression of cell surface exhaustion markers (8–14). This immune system dysfunction impacts the ability of CD8+ T cells to suppress viral replication (15–18). For normal progressors, the clinical latency phase lasts an estimated 10 years and ends with an eventual decline in their CD4+ T cell count which results in the inability to fight opportunistic infections, marking their transition to AIDS (3). While most HIV-1 patients lose full immune function within 8-10 years after infection, a small subset deemed HIV-1 controllers display spontaneous and robust control of HIV-1 replication and subsequently maintain significantly slower disease progression and low viral loads without therapy (3, 19). Reflective of their low viral loads, these individuals retain a normal CD4+ T cell count without drug intervention, which consequently extends their clinical latency phase and improves their life expectancy (20–23). Understanding the molecular mechanisms that permit HIV-1 controllers to suppress viral replication can aid in the development of novel HIV-1 immunotherapeutics or vaccines.

HIV-1 infection results in a progressive cascade of immunological events that lead to chronic immune activation and inflammation, which remain persistent due to the ability of HIV-1 to evade immune detection and elimination. This is mainly caused by the large-scale depletion of CD4+ T cells that regulate the adaptive immune response, and the inability of ART to eliminate latently infected cells (24–29). Under normal conditions, activated T cells respond to infection through the secretion of cytokines, clonal expansion, proliferation, and degranulation. Upon activation, T cells also up-regulate immune checkpoint receptors on their surface which can both negatively regulate the immune response to infection and maintain self-tolerance (30). During acute infection, immune checkpoints are part of a homeostatic mechanism which attenuates the immune response, creating a negative feedback loop to abrogate T cell activation as the infection is resolved. This mechanism prevents tissue damage from chronic inflammation caused by continued T cell activation (31, 32). However, during chronic infection, T cell activation persists and the immune system must adapt to minimize tissue damage while still maintaining control of infection. To mitigate immunopathology, the immune system has evolved mechanisms to progressively temper the immune response during chronic infection. Over time this leads to a state of hypo-responsiveness and eventual immunological tolerance of the virus (33, 34). T cell exhaustion is one such mechanism evolved to temper the immune response in which effector T cells upregulate inhibitory immune checkpoint receptors (ICR) resulting in hierarchical loss of T cell function (9, 13, 14, 35). T cell exhaustion represents a spectrum of predictable, progressively impaired effector functions that are a direct consequence of chronic infection (13, 33, 36).

T cell exhaustion impacts the ability of CD8+ T cells to suppress viral replication and has been characterized by the co-expression of multiple ICRs (15–18, 31). ICRs, when engaged with their corresponding ligand, negatively regulate T cell activation, and restrict the effective control of infection (37–39). Programmed cell death 1 (PD-1) is one such ICR whose expression is up-regulated on CD8+ T cells in both chronic viral infections and cancer (40, 41). Initial up-regulation of PD-1 on CD8+ T cells during the acute phase of infection does not restrict cytotoxic activity (42). However, during chronic infection, increased PD-1 expression on CD8+ T cells yields an exhausted phenotype and subsequent failure to control HIV viral replication (8, 10, 12, 40). It was demonstrated that ART was not sufficient to restore the polyfunctional, proliferative, or cytotoxic capabilities of dysfunctional HIV-specific CD8+ T cells (43). In addition, it was demonstrated that the length of time between initial infection and initiation of ART impacts the ability of treatment to successfully reduce markers of T cell exhaustion (44, 45). T-cell immunoglobulin and mucin domain-3 (Tim-3) is another inhibitory immune checkpoint receptor implicated in T cell exhaustion, and is known to play a role in the reduction of IFN-ƴ mediated inflammation (46). Tim-3 has been observed to be upregulated during chronic infection with HIV and its expression correlates to clinical parameters of disease progression (47). The expression of Tim-3 has also been shown to be upregulated despite treatment with ART (48). In addition, the co-expression of both PD-1 and Tim-3 has been observed to be upregulated in chronic infection and represents the most severe exhausted phenotype (41, 47, 49, 50). Other ICRs upregulated in T cell exhaustion include cytotoxic lymphocyte antigen-4 (CTLA4), T-cell immune receptor with Ig and ITIM domains (TIGIT), and lymphocyte-activation gene 3 (LAG3) (18). We hypothesize that efficient control of viral replication becomes compromised by CD8+ T cell exhaustion.

Our previous work identified that the ability of some people living with HIV (PLWH) to maintain a low viral load in the absence of therapy was accompanied by the ability of PBMCs from these individuals to suppress replication of an HIV molecular clone ex *vivo*. This *ex vivo* control was shown to be mediated by CD8+ T-cells. Further, CD8+ T-cells in these controllers maintained levels of activation markers and the exhaustion marker PD-1 equivalent to uninfected individuals, unlike PLWH without a history of control who exhibited the expected increase in activation and PD-1 (51).

Despite their initial inherent ability to suppress viral replication in the absence of ART, mechanisms that are poorly understood cause HIV-1 controllers to eventually lose the ability to control viral replication as measured by an increase in HIV plasma viral load. We captured this phenomenon in four HIV-1 controllers (designated LVL post-control) who were consequently placed on ART. As expected, we observed a subsequent loss of *ex vivo* resistance to HIV infection in PBMCs from these donors. This gave us the unique opportunity to observe real time changes associated with HIV-1 control by examining longitudinal samples from these donors to determine whether the loss of control is associated with CD8+ T cell exhaustion. We hypothesized that changes in the activation and exhaustion state of T-cells in our LVL cohort would correlate with both *in vivo* and *ex vivo* control and that addressing exhaustion via immune checkpoint blockade might support prolonged control of infection. We therefore characterized the progressive changes in a broad panel of CD8+ T cell exhaustion markers using our cohort of HIV-1 controllers to determine if changes in activation and exhaustion correspond to the loss of HIV-1 control and tested the ability of immune checkpoint blockade (ICB) targeting exhaustion markers to restore *ex vivo* control in LVLs who recently lost control.

## RESULTS

### HIV controllers have altered profiles of CD8+ T-cell exhaustion

We hypothesized that the ability of HIV controllers with low viral loads (LVLs) to control HIV replication in the absence of therapy may be related to effective control of CD8+ T-cell exhaustion. To examine this, we performed flow cytometry on an expanded set of PBMCs from LVLs to characterize their exhaustion and activation phenotypes. Blood was drawn from HIV-negative individuals (ND or normal donor, n=5), LVLs (n=5), and HIV-1 normal progressors with high viral loads and no ART treatment (HVLs) (n=4) during scheduled clinic visits (Table 1). PBMCs isolated from blood samples were stained and examined by flow cytometry. After gating for single, live CD3+ cells, CD8+ T cells were first examined for the expression of activation markers CD38 and HLA-DR (Fig 1). LVLs exhibited an activation profile similar to NDs, with significantly lower enrichment of CD38+ (Fig 1D) and HLA-DR+ (Fig 1E) CD8+ T cells compared to HVLs. LVLs also exhibited significantly less CD38+/HLA-DR-(Fig 1G) and CD38+/HLA-DR+ (Fig 1I) CD8+ T cells than HVLs, again similar to ND. The high level of activation markers expressed on CD8+ T cells in HVLs confirmed previous reports that chronic HIV infection is associated with immune activation (52–58). This data also indicates that LVLs, despite maintaining detectable viraemia, maintain low levels of immune activation comparable to HIV uninfected individuals.

**Figure 1:**
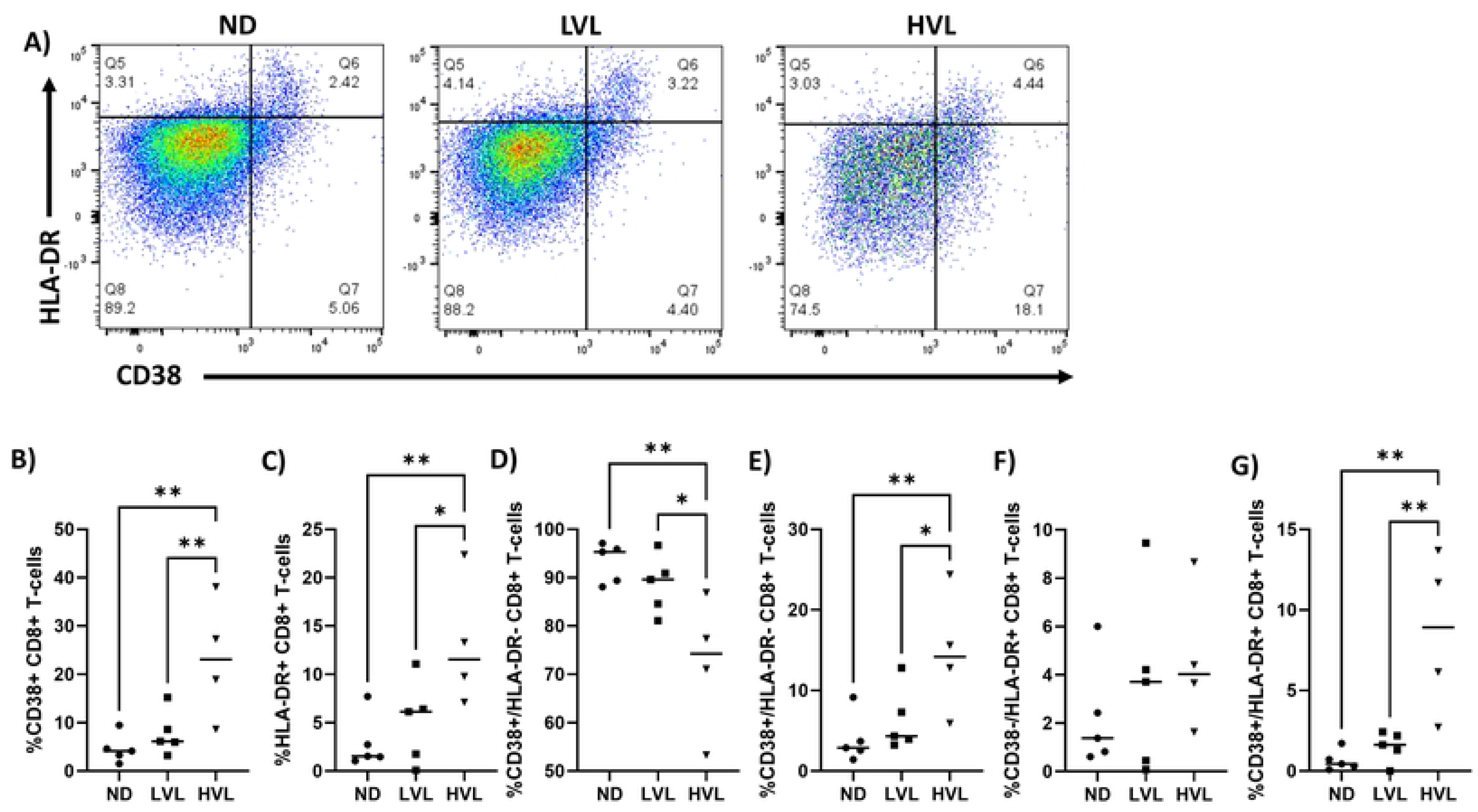
HIV-1 controllers maintain low levels of CD8+ T-cell activation. PBMCs were isolated from the blood of HIV negative subjects (ND), HIV controllers off therapy (LVL) and HIV standard progressors off therapy (HVL). **A)** Flow cytometry was performed and live (Zombie Yellow-), CD8+ T-cells (CD3+/CD8+) were analyzed for the expression of CD38 and HLA-DR. The percentage of CD8+ T-cells positive for each marker was determined for each donor and graphed: **B)** all CD38 positive, **C)** all HLA-DR positive, **D-G)** CD8+ T-cells with the given combination of CD38 and HLA-DR. Normality was evaluated by Kolmogorov-Smirnov test and one-way ANOVA was performed to determine significance. * p≤0.05, ** p≤0.01

**Table 1:** Donor characteristics at study enrollment.

| Classification | Subject ID | VL<br>[copies/ml] | CD4<br>[cells/mm3] | CD8<br>[cells/mm3] | ART | Therapy |
| --- | --- | --- | --- | --- | --- | --- |
| HVL | ADV7140 | <20 | 497 | 593 | ON | Truvada/Tivicay |
|  | BRW1143 | <20 | 1135 | 828 | ON | Truvada/Tivicay |
|  | XTD8730 | 30 | 251 | 764 | ON | Tivicay/Reyataz/Norvir |
|  | KLU1328 | <20 | 337 | 794 | ON | Descovy/Tivicay |
|  | UNJ7200 | <20 | 317 | 517 | ON | Complera |
|  | KMJ1960 | 30,000 | 145 | 1367 | off | - |
|  | WBZ1300 | 24,000 | 170 | 1206 | off | - |
|  | ORL5590 | 2,880,820 | 62 | 344 | off | - |
|  | ARF1454 | 35,325 | 39 | 509 | off | - |
| LVL | FJG8070 | 250 | 1070 | 1434 | off | - |
|  | FWU1270 | 100 | 688 | ND | off | - |
|  | HCQ6670 | 80 | 488 | 506 | off | - |
|  | MPY1313 | 1,550 | 473 | 756 | off | - |
|  | NIM1164 | 750 | 723 | 987 | off | - |
|  | SRS5930 | 230 | 699 | 792 | off | - |
|  | VQY4910 | 750 | 723 | 987 | off | - |
|  | AEM9650 | 1630 | 1001 | 2081 | off | - |
| Normal Donor | BTS1096 | - | 1190 | 358 | - | - |
|  | CHT3368 | - | 360 | 646 | - | - |
|  | HFK1114 | - | 840 | 597 | - | - |
|  | ENS1170 | - | - | - | - | - |
|  | VIQ1422 | - | - | - | - | - |
| Elite Controller | EXT1011 | <20 | 542 | 498 | ON | Yes |

We next examined the expression of exhaustion markers PD-1 and TIM3, on CD8+ T cells from NDs, LVLs, and HVLs (Fig. 2). When considering PD-1 and TIM3 expression alone, CD8+ T cells from LVLs again displayed a similar phenotype to NDs (Fig. 2D, E). When considering PD-1 and TIM3 coexpression, LVLs exhibited a significantly lower enrichment of PD1-/TIM3+ (Fig. 2G), PD1+/TIM3-(Fig. 2H), and PD1+/TIM3+ (Fig. 2I) CD8+ T cells compared to HVL. Similar to the observed activation profiles, no significant differences in exhaustion marker expression were observed between ND and LVL (Fig 2D-I).

**Figure 2:**
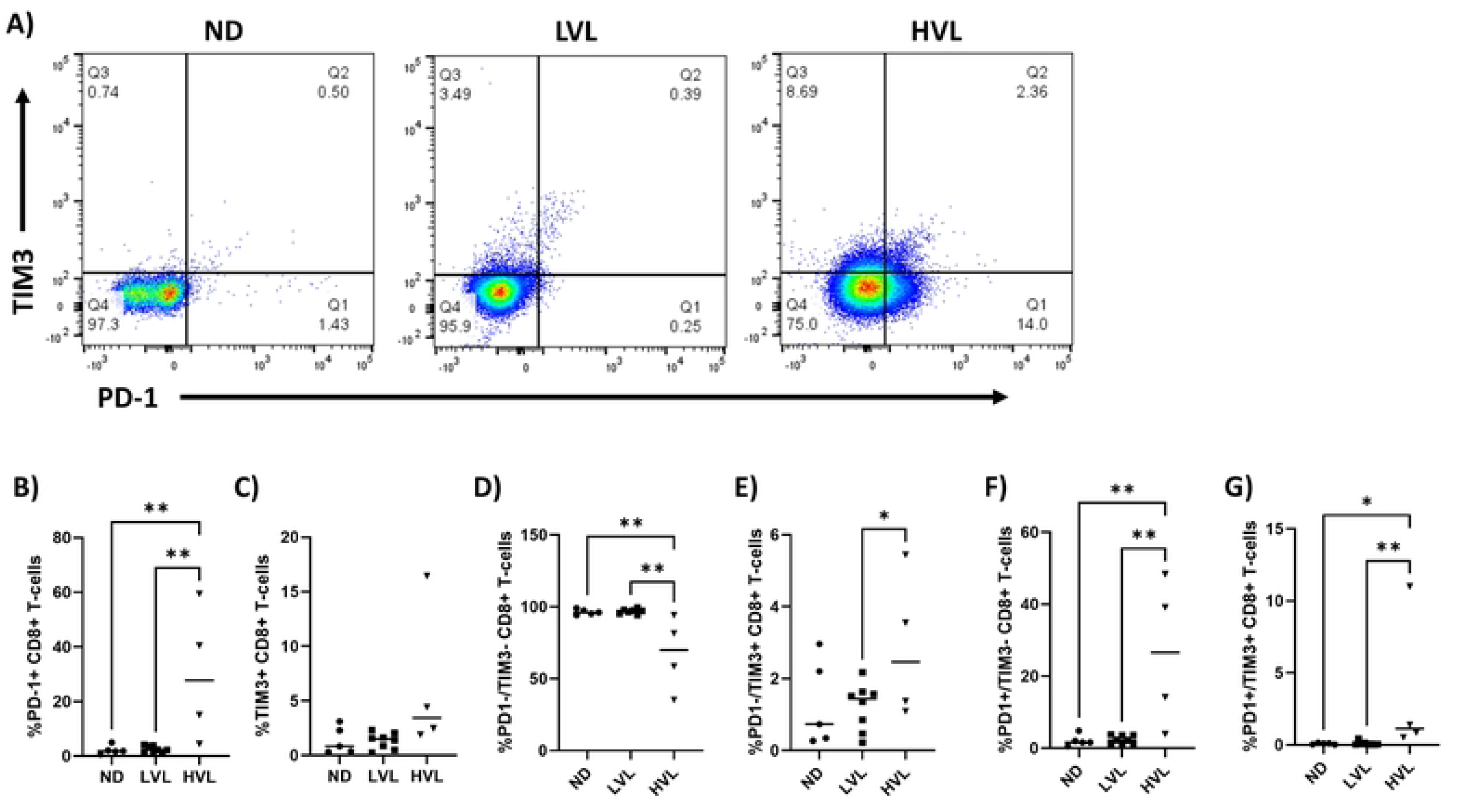
HIV-1 controllers maintain low levels of PD-1 and TIM3 on CD8+ T-cells. PBMCs were isolated from the blood of HIV negative subjects (ND), HIV controllers off therapy (LVL) and HIV standard progressors off therapy (HVL). **A)** Flow cytometry was performed and live (Zombie Yellow-), CD8+ T-cells (CD3+/CD8+) were analyzed for the expression of PD-1 and TIM3. The percentage of positive CD8+ T-cells was determined for each donor and graphed: **B)** all PD-1 positive, **C)** all TIM3 positive, **D-G)** CD8+ T-cells with the given combination of PD-1 and TIM3. Normality was evaluated by Kolmogorov-Smirnov test and one-way ANOVA was performed to determine significance. * p≤0.05, ** p≤0.01

To expand our examination of exhaustion markers, we determined the levels of CTLA-4, LAG3, and TIGIT in CD8+ T cell populations of the same three groups (Fig. 3). LVLs exhibited a significantly higher enrichment of CTLA-4+ CD8+ T cells compared to HVLs (Fig. 3A). This may be explained by the fact that the HIV-1 nef protein, which should be highly expressed in HVL donors off ART, has been shown to downregulate CTLA-4 (59). A similar trend was observed for LAG3 (Fig. 3B), with the opposite trend displayed for TIGIT (Fig. 3C). Interestingly, the similarities in activation and exhaustion profiles seen in LVLs and NDs were not observed in this analysis as CTLA-4, LAG3, and TIGIT expression were significantly different between the groups. Taken together, the data in figures 2 and 3 support the conclusion that LVL controllers display a unique exhaustion phenotype compared to HIV-1 normal progressors and HIV-uninfected individuals. Specifically, they maintain low levels of TIM3+, PD-1+, and TIGIT+ CD8+ T cells, while showing increased enrichment of CTLA-4+ and LAG3+ CD8+ T cells.

**Figure 3:**
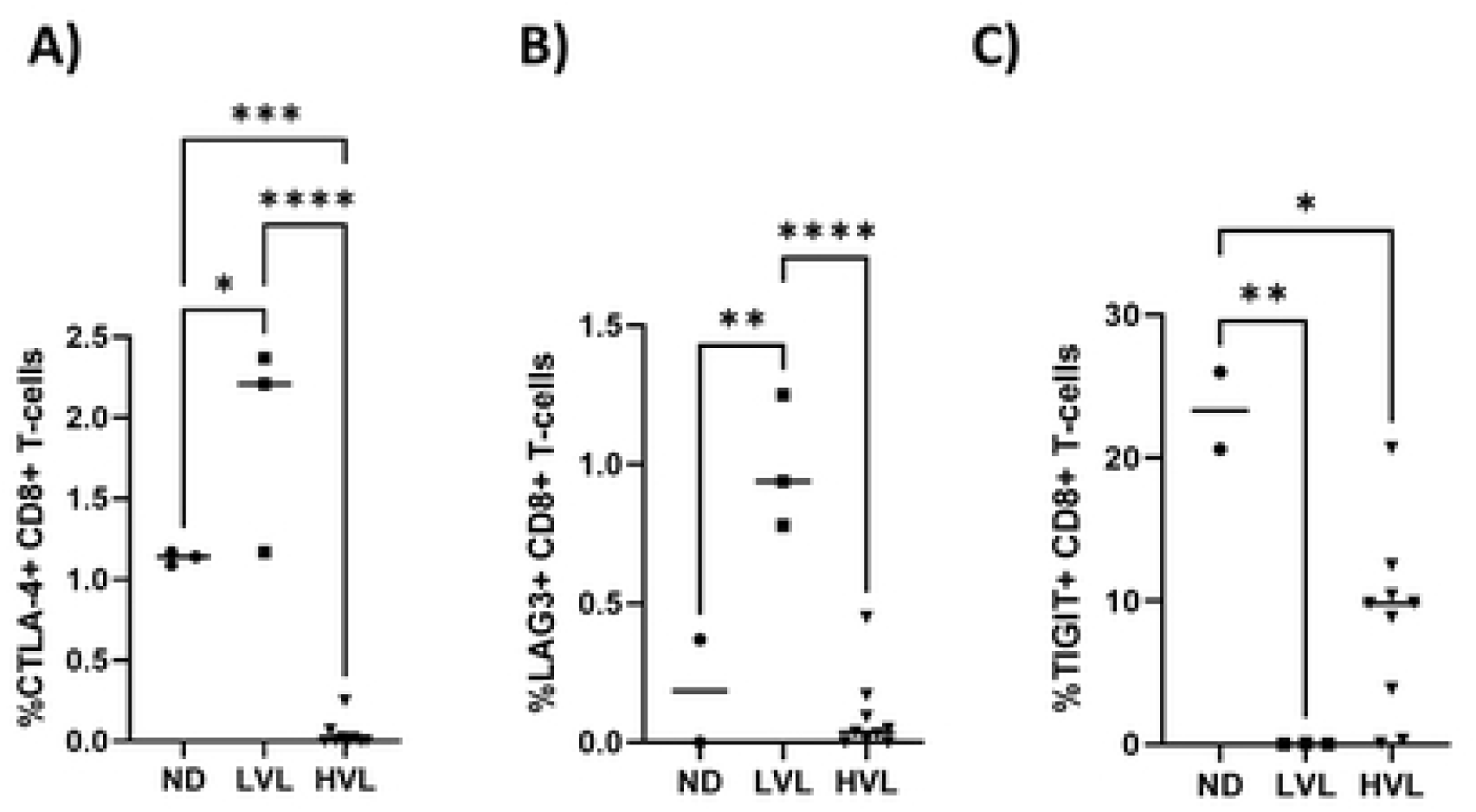
Levels of CTLA-4, LAG3 and TIGIT on CD8+ T-cells of normal donors, LVLs and HVLs. PBMCs were isolated from the blood of HIV negative subjects (ND), HIV controllers off therapy (LVL) and HIV standard progressors off therapy (HVL). Flow cytometry was performed and live (Zombie Yellow-), CD8+ T-cells (CD3+/CD8+) were analyzed for the expression of **A)** CTLA-4, **B)** LAG3, or **C)** TIGIT. Graphed is the percentage of CD8+ T-cells positive for each marker. Normality was evaluated by Kolmogorov-Smirnov test and one-way ANOVA was performed to determine significance. * p≤0.05, ** p≤0.01

### Loss of control is associated with increased CD8+ T cell exhaustion

Over the course of this study, four LVL participants experienced an increase in viral load and were placed on ART, which successfully reduced their viral load to undetectable levels. Our previous study demonstrated that PBMCs from LVLs exposed to HIV-1 *ex vivo* did not support viral replication due to the presence of CD8+ T-cells and that this ability was lost in three of the four LVLs after losing control and being placed on ART. We examined the levels of activation (CD38/HLA-DR) and exhaustion (PD-1/TIM3) on CD8+ T-cells from PBMCs of these LVL post control individuals after being placed on therapy. These levels were then compared to LVLs during periods of successful control (Fig. 4). No significant differences were observed in the levels of CD8+ T-cell activation (Fig. 4A-F). However, the levels of exhaustion as measured by PD-1 and TIM3 expression increased both for each marker individually and in combination (Fig. 4G-L) in CD8+ T cells from LVLs who had lost control. Whether CD8+ T cell exhaustion was the cause, or a result of the loss of control is unclear. However, the observed association between loss of control and CD8+ T cell exhaustion supports our hypothesis that the ability of LVLs to control HIV replication is related to effective control of CD8+ T-cell exhaustion.

**Figure 4:**
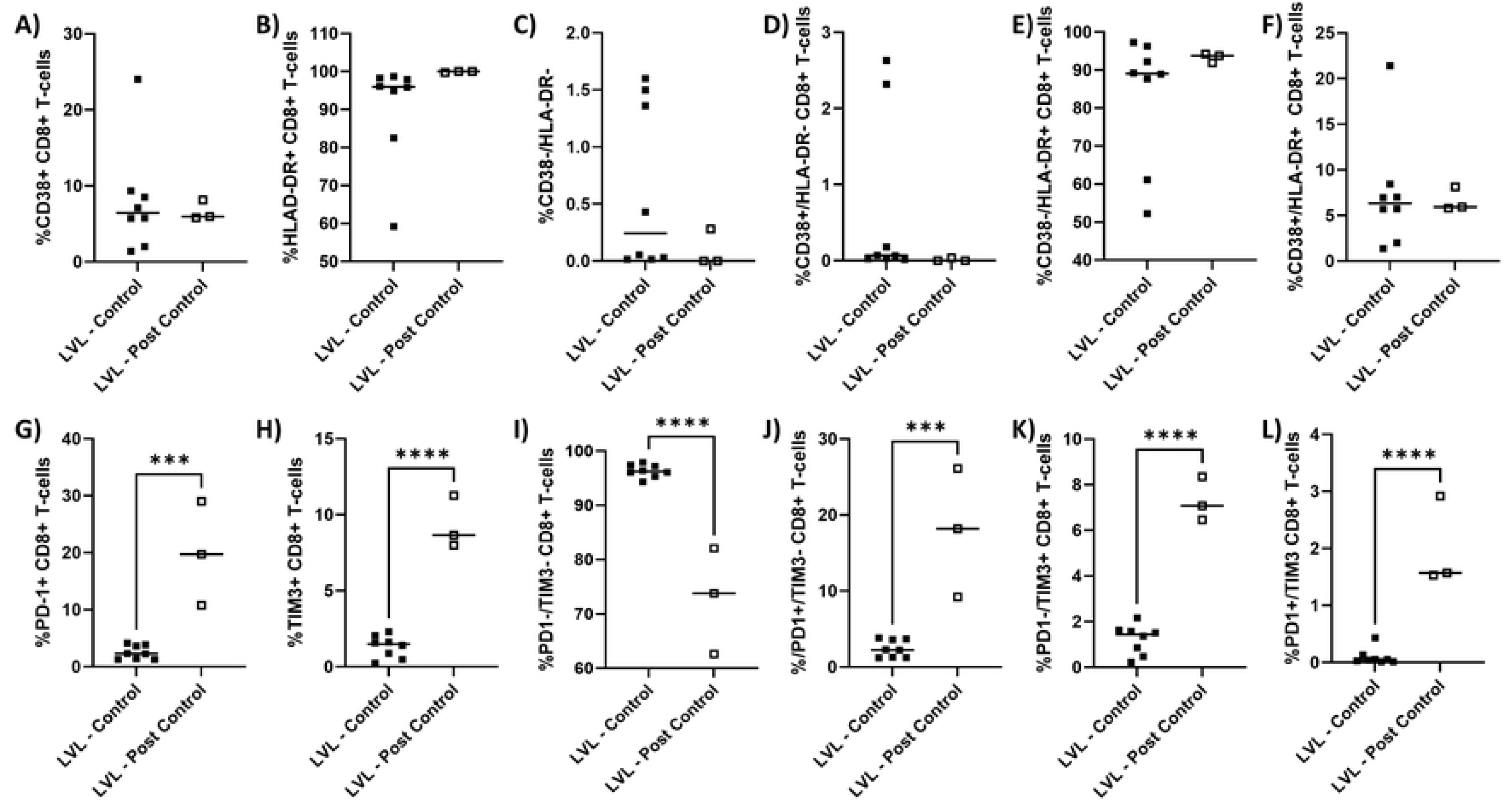
Comparison of CD8+ T-cell activation and exhaustion states in LVLs versus LVLs post control. PBMCs from LVL controllers (LVL – Control) or LVL post control (LVL – Post Control) were evaluated by flow cytometry. The percentage of **A-F)** CD38 and HLA-DR and **G-L)** PD-1 and TIM3 on live, CD3+/CD8+ T-cells was determined. Normality was evaluated by Kolmogorov-Smirnov test and one-way ANOVA was performed to determine significance. *** p≤0.0005, **** p≤0.0001

### Multivariate analysis reveals heterogenous populations of T cells expressing various combinations of activation and exhaustion markers

Complex immune states like T cell exhaustion are related to the simultaneous expression of multiple markers. Traditionally, these states are interrogated by looking at combinations of specific markers, as above, or assessing polyfunctionality by looking at the number of co-expressed markers. To evaluate the importance of the expression of individual markers in a qualitative manner independent of gating choices, we performed clustering analysis using flow cytometry data from NDs, LVLs, LVLs post control and HVLs on and off ART (Fig 5). Populations identified as live, CD3+ T-cells were clustered using the FlowSOM plugin for FlowJo v10. This clustering revealed multiple populations of CD4+ and CD8+ T-cells displaying unique combinations of activation and exhaustion markers (Fig. 5A). These populations were color coded and designated 1-10. To confirm these clusters and determine whether specific clusters are overrepresented in any group, we performed dimensional reduction using t-Distributed Stochastic Neighbor Embedding (tSNE) (Fig. 5B). The tSNE plot confirmed the clustering analysis and as expected, revealed that some of the ten clusters identified might be further broken down into sub clusters (i.e. cluster 7 or 8 which both occupy multiple distinct regions on the tSNE plot). By displaying cells from each donor classification alone, we were able to determine which clusters and regions were shared or unique to each class of donor (Fig. 5C-G). Normal donors were characterized by the presence of CD8+ T-cells (cluster 7) or CD4+ T-cells (CD8-cells, cluster 10) with low levels of CD38 and HLA-DR. HVLs displayed multiple populations of HLA-DR+ and CD38+ CD8+ T-cells (cluster 2) along with some smaller populations with low expression of HLA-DR and CD38 (cluster 7). LVLs who maintained control showed a profile with similarities to both NDs and HVLs, but resembled ND more closely. Interestingly, LVLs who maintained control also possessed unique populations not seen in NDs or HVLs (labeled 1* and 8* as sub-populations of clusters 1 and 8 respectively, indicated by outline in panel E). As the LVLs who lost the ability to control were placed on ART, we also analyzed HVLs who were suppressed on therapy to account for any contribution of ART. Interestingly, LVLs who lost control and were placed on ART displayed profiles similar to HVLs on ART, with a unique region shared between them (labeled 4* indicated by outline in panel F and G) representing CD8-cells with high levels of activation markers. LVLs who lost control also displayed a unique population not seen in any other group (labeled 5* in panel F). To further characterize the phenotypes of T cells in the regions unique to LVLs, we displayed the expression of CD8, CD38, HLA-DR, PD-1, TIM3, TIGIT, CTLA4 and LAG3 as a heatmap projected onto the tSNE plots (Fig 6). This showed that population 1*, a population unique to LVLs who maintained control, was composed of CD8+ T-cells with intermediate HLA-DR expression, and very low expression of PD-1, CTLA-4 and TIM3. Population 8*, another population unique to LVLs with control, had mixed levels of HLA-DR and CD38, but expressed higher levels of exhaustion markers including CTLA-4 and LAG3. LVLs who lost control had profiles similar to HVLs on ART in that they had high levels of activation, but also included unique populations not seen in either normal donors or HVLs off ART (populations 4* and 5*). Heatmap analysis revealed these populations to likely be CD4+ T cells (CD3+, CD8-) with higher levels of HLA-DR and CD38, although variability was still high within the population (i.e. HLA-DR). Population 4* was also present in HVLs on suppressive therapy, but population 5*, which express TIM3, TIGIT, and CTLA4, but not PD-1, was absent in these HVLs on therapy. Taken together, these data support the idea that LVLs maintain a low level of CD8+ T cell exhaustion as characterized by the low abundance of PD-1+ and TIM3+ CD8+ T-cells, and upon loss of control and subsequent ART treatment, display activation and exhaustion phenotypes similar to HVLs on ART.

**Figure 5:**
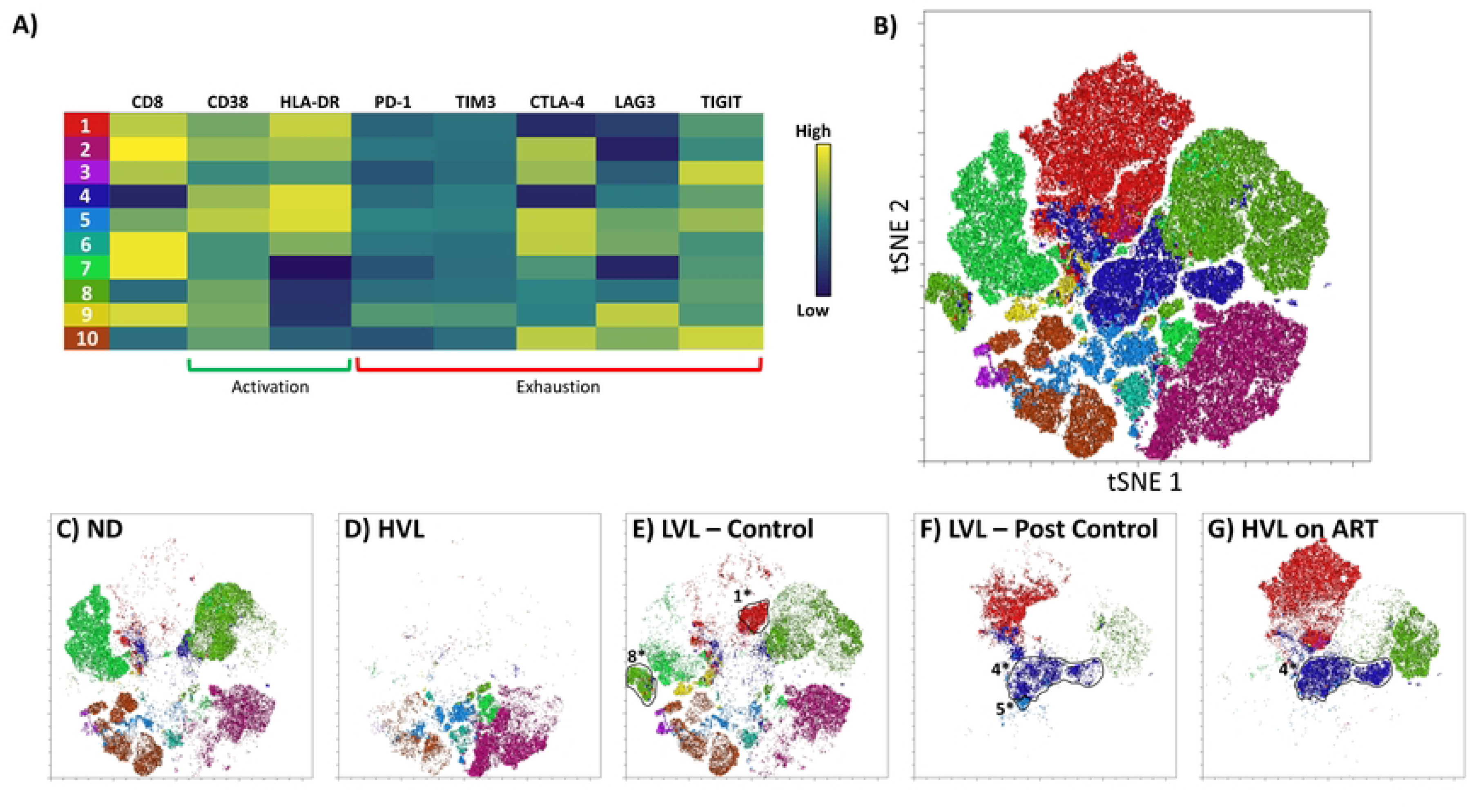
Clustering and tSNE dimensionality reduction of flow cytometry profiles of CD8+ T-cells from different subject groups. Flow cytometry was performed on PBMC from normal donors, HVL, LVL and LVL post control using the Zombie Yellow live/dead stain and antibodies for CD3, TIM3, CD38, TIGIT, PD1, CTLA4, HLA-DR, LAG3, and CD8. **A)** Cells identified as live (Zombie Yellow-) T-cells (CD3+) were clustered using the FlowSOM plugin in FlowJo. **B)** tSNE analysis and plotting was performed on the same live T-cell populations using the associated function in FlowJo. Final plot was colored according to the clusters identified in panel A. Cells from human subjects corresponding to **C)** HIV negative (ND), **D)** HVL, **E)** LVL, **F)** LVL post control, and **G)** HVL on ART were mapped back onto the tSNE plot. Circled populations are those unique to either LVLs, LVLs post control, or HVLs on ART and are labelled with an identifying number that corresponds to the cluster (A) of which they are a sub-population.

**Figure 6:**
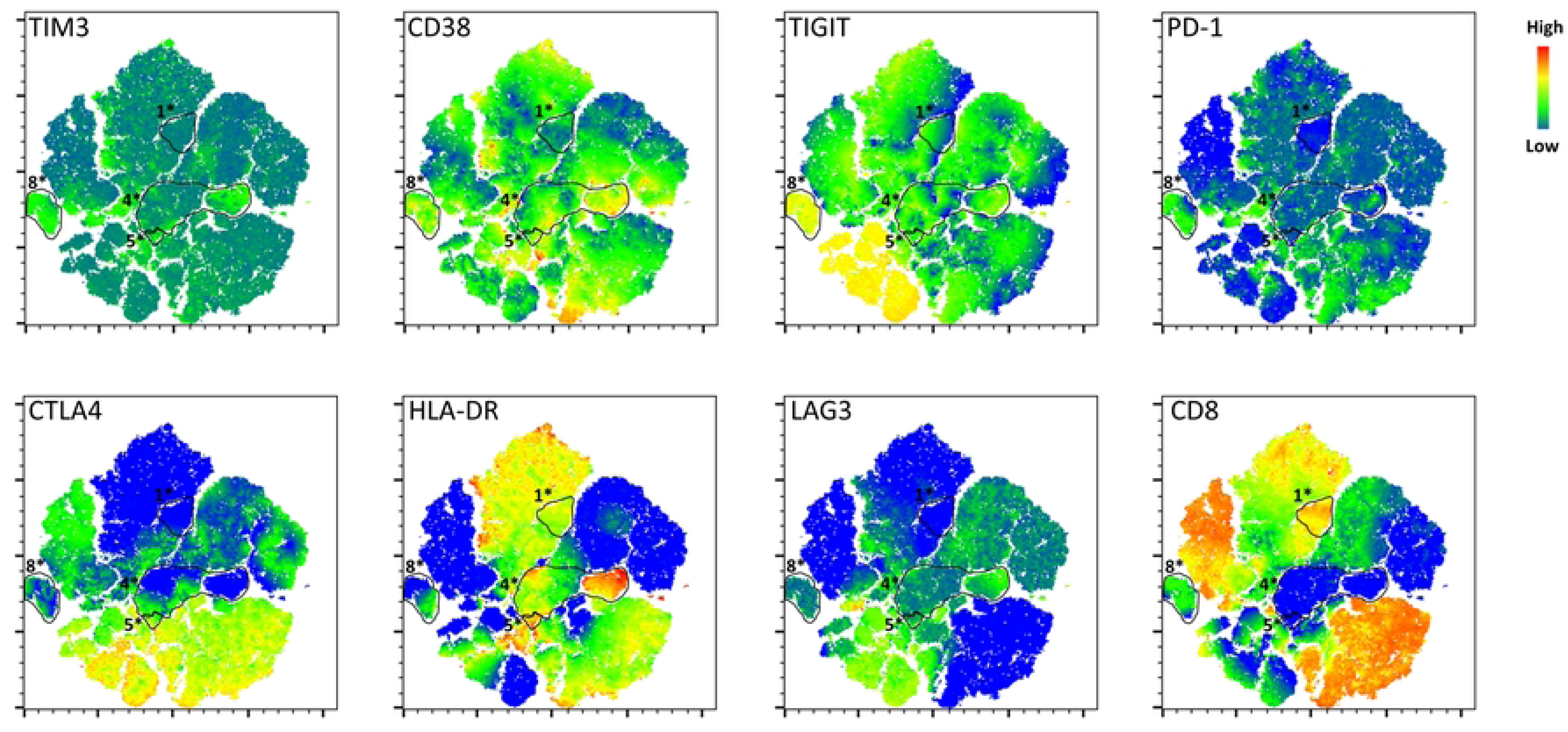
Expression of individual markers by different populations as defined by tSNE. The intensity of staining for each individual marker was mapped back onto the tSNE plot generated in figure 5. Heatmap indicates the relative expression levels of each marker from low (blue) to high (red). The circled and labeled populations are those that were unique to LVLs (1* and 8*), LVLs post control (4* and 5*), and HVLs on ART (4*) and are identified according to the cluster they were assigned to in figure 5.

### PD1/TIM3 levels increase over time on therapy in LVLs who lost control

Previous studies have suggested that treatment with ART can reduce PD-1 expression on CD8+ T cells (8, 10, 12, 60–63), while some suggest that Tim-3 expression persists despite treatment with ART (48, 64, 65). However, exhaustion marker expression patterns in controllers post-control treated with ART remain unknown. Therefore, we sought to determine whether the increase in Tim-3 and PD-1 expression on CD8+ T cells observed during the loss of *in vivo* viral control (Fig. 4) persisted once these patients were ART suppressed. Once ART is initiated, the rate of viral decay varies across individuals, but typically occurs in three phases (66–68). Therefore, we pooled longitudinal donor data according to three timeframes corresponding to these phases; <180 days, 181-365 days, and 366+ days. To determine the effects of ART on T cell exhaustion, we compared the pooled longitudinal T cell exhaustion flow cytometry data from LVLs post control and compared it to ART naïve LVLs during periods of *in vivo* and *ex vivo* viral control and to HVLs suppressed on ART (Fig 7A-D). Significant increases in the percentage of PD1+/TIM3-and PD1+/TIM3+ CD8+ T-cells were observed after, or just prior to one year on therapy, respectively, suggesting populations of exhausted T cells persist despite ART (Fig. 7B, D). HVLs on therapy also showed increased levels of exhausted CD8+ T cells with significantly more PD1-/TIM3+ cells compared to LVLs with control, suggesting that ART does not reverse T cell exhaustion states.

**Figure 7:**
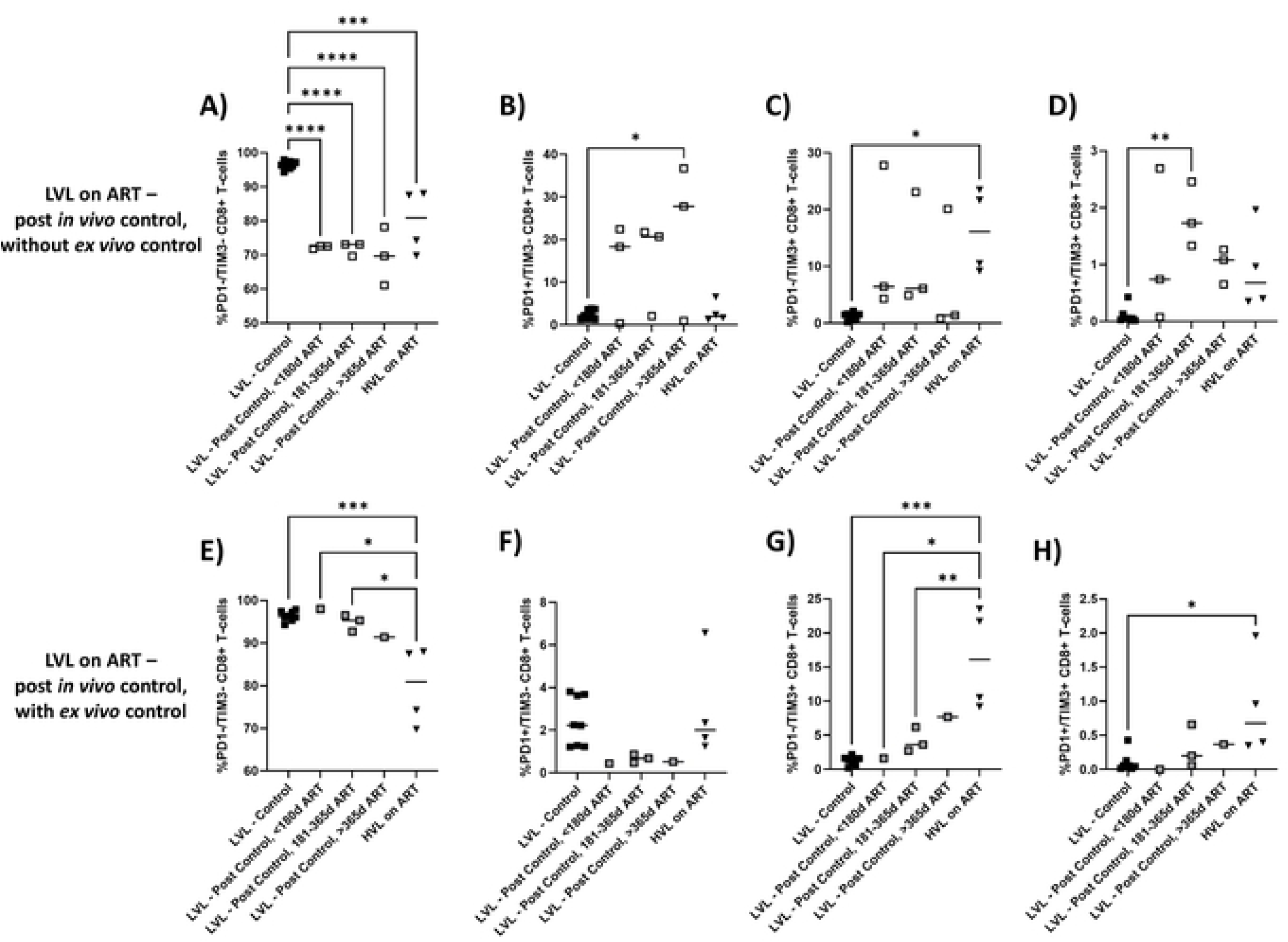
Changes in PD1 and TIM3 exhaustion markers in CD8+ T-cells in LVL and LVL post control after beginning anti-retroviral therapy. PBMCs from LVLs post control (**A-D**) and LVL who maintained ex vivo control (**E-H**) were analyzed by flow cytometry for the expression of PD1 and TIM3 on CD8+ T-cells. Expression of different combinations of these two markers is shown for samples drawn from patients at different times post initiation of ART (<180 days, 181-365 days and 366+ days). Levels of expression are compared to LVL controllers pre-therapy and to HVL on therapy.

Since the expression patterns of PD-1 and Tim-3 on CD8+ T cells has been suggested to identify different levels of T cell exhaustion with functional implications, we next aimed to determine whether the exhaustion profile observed in ART suppressed LVL donors who maintained *ex vivo* control changed over time on therapy. To evaluate this, we once again pooled exhaustion data from donors who continued to maintain *ex vivo* control on ART and grouped them according to three timeframes <180 days on ART, 181-365 days on ART, and 366+ days on ART. There was no significant difference in the levels of Tim-3-/ PD-1+, Tim-3+/ PD-1-, nor Tim-3+/ PD-1+ CD8+ T cells across all time points for ART suppressed LVLs who maintained *ex vivo* control in comparison to ART naïve LVLs who could control (Fig 7 F-H). These results further display the association between low CD8+ T cell exhaustion levels and the ability to control viral replication both *in vivo* and *ex vivo*.

### Immune checkpoint blockade restores *in vitro* control in pure cultures of T-cells

Since we observed an association between CD8+ T cell exhaustion and the loss of control in our cohort of LVL donors, we hypothesized that immune checkpoint blockade (ICB) targeting exhaustion markers could potentially restore *ex vivo* control in these donors. To test this hypothesis, and determine whether CD8+ T cell exhaustion mediates the loss of *ex vivo* viral control, activated CD8+ T cells were treated with anti-PD-1 and anti-Tim-3 ICB antibodies and evaluated for their ability to suppress HIV replication *in vitro* (69, 70). CD8+ T cells and autologous CD4+ T cells were isolated from LVLs post control using a magnetic bead based negative isolation. CD8+ and CD4+ T cell isolates were induced with PHA and IL-2 for 48 hours and co-cultured at a 1:1 ratio in the presence or absence of ICB. All cells were infected with HIV NL4-3and supernatant was collected and lysed every 48 hours post infection. HIV p24 concentrations were measured via ELISA as a readout of viral replication. When anti-PD-1 (Fig 8A) or anti-Tim-3 (Fig 8B) ICB were titrated, there was no appreciable decrease in supernatant p24.

**Figure 8:**
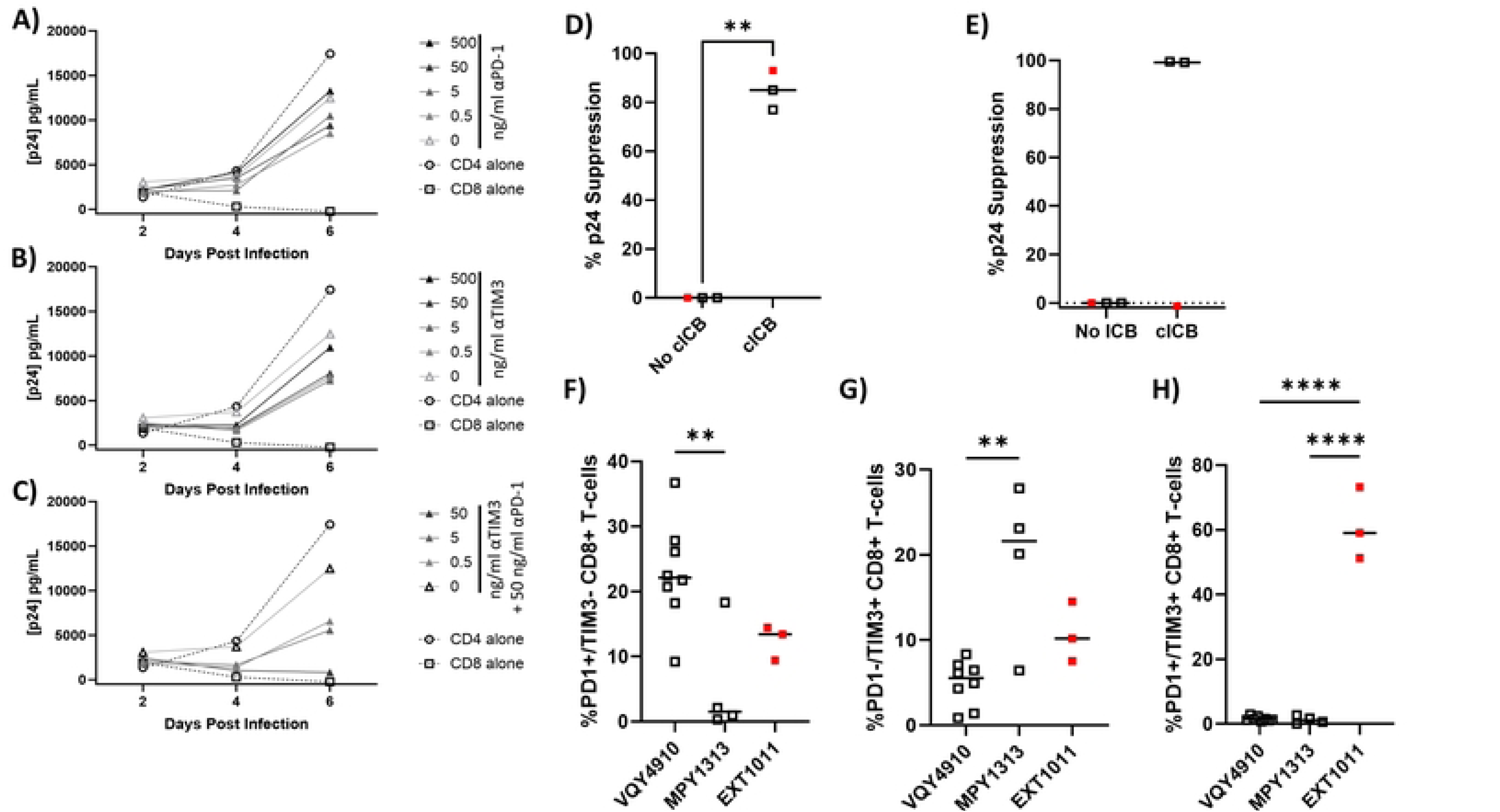
Combination immune checkpoint blockade can restore *ex vivo* control of viral replication in LVLs post control and CD8+ T cell exhaustion profiles are associated with cICB responsiveness. CD4+ and CD8+ T-cells were isolated from PBMC of LVL post control. CD4+ and CD8+ T-cells were activated by PHA and IL-2 and cultured alone or in a 1:1 combination and infected with NL4-3. Cultures were maintained in the given concentration of **A)** anti-PD-1 antibody alone, **B)** anti-TIM3 antibody alone, or **C)** anti-TIM3 antibody with 50 ng/ml anti-PD-1. Supernatant levels of p24 were determined by Elisa on days 2, 4 and 6 post infection. Ability of 50 ng/ml of anti-PD-1 and 50 ng/ml of anti-TIM3 to restore suppression of viral replication in cultures of multiple donors’ **D)** purified CD4+ and CD8+ T-cells or **E)** total PBMCs was determined by computing the percentage of suppression at peak p24 levels as compared to mock treated cultures. LVL post control on therapy are indicated as black symbols and an elite controller on therapy is indicated as a red symbol ** p≤0.01. PBMCs from blood draws performed at multiple time points following the beginning of ART in LVL post control (VQY4910 and MPY1313) and an elite controller on therapy (EXT1011) were examined by flow cytometry for the percentage of **F)** PD1+/TIM3-, **G)** PD1-/TIM3+ and **H)** PD1+/TIM3+ CD8+ T-cells.

To test the combination of ICB (cICB), anti-Tim-3 was titrated in the presence of 50 ng/mL anti-PD-1 ICB (Fig 8C). The lowest dose of anti-TIM-3 (0.5 ng/mL) in the presence of anti-PD-1 resulted in an approximate 50% suppression of viral replication compared to 50 ng/mL anti-PD-1 alone (Fig 8C, light gray triangles). Strong suppression of viral replication was restored in the 50 ng/mL anti-TIM3 and 50 ng/mL PD-1 dosing (Fig 8C, black triangles). When additional ART-treated LVL donors (n=3) who could not suppress viral replication *ex vivo* were pre-treated with anti-Tim-3 and anti-PD-1 cICB, suppression of viral replication was restored (Fig 8D), further supporting the hypothesis that CD8+ T cell exhaustion restricts effective control of viral replication in LVLs who lose control.

### Response to immune checkpoint blockade is associated with CD8+ T cell exhaustion levels in PBMC cultures

CD8+ T cell functionality is dependent on a multitudinous array of complex and integrated signals from the surrounding microenvironment, including a variety of signals that have both stimulatory and inhibitory effects on T cell activation (71–73). To better capture the effects of these important signaling interactions, we aimed to determine whether the restored CD8+ T cell functionality gained with the treatment of cICB in CD4+ and CD8+ T cell co-cultures would also exist in the context of total PBMCs. To examine this, PBMCs from LVLs post control and an elite controller on therapy were induced with PHA and IL-2 for 48 hrs and cultured in the presence or absence of 50 ng/mL anti-PD-1 and 50 ng/mL anti-Tim-3 cICB for 24 hrs. PBMCs were either infected with HIV NL4-3 virus or cultured in media alone, the supernatant was collected and lysed every 48 hours post infection and HIV p24 concentrations were measured via ELISA. In the absence of cICB, no donors displayed the ability to suppress viral replication *in vitro* as measured by the inability to suppress production of viral p24 (Fig 8E). However, there was a dichotomous response of PBMCs to cICB across donors. Two of the donors (VQY4910 and MPY1313) showed restoration of p24 suppression in PBMCs upon cICB treatment. However, the elite controller (EXT1011), despite showing restoration of control when using isolated CD4+ and CD8+ T cells, did not show restoration of control in PBMC cultures (Fig 8E).

To understand the differential response to cICB, we directly compared the exhaustion levels of VQY4910 and MPY1313, the cICB responsive donors, and EXT1011, the cICB non-responsive donor, via flow cytometry analyzing expression of Tim-3 and PD-1 on CD8+ T cells. Heterogeneity was seen within cICB responsive donors, as VQY4910 had a significantly higher percentage of PD-1+/Tim-3-CD8+ T cells (Fig. 8F) and a significantly lower percentage of PD-1-/TIM3+ CD8+ T cells compared to MPY1313 (Fig 8G). Assessing the more severely exhausted phenotype, we observed a significantly higher enrichment of PD-1+/Tim-3+ CD8+ T cells from EXT1011 than both MPY1313 and VQY4910 (Fig. 8H). When we evaluated other clinical markers of disease progression, we noted that all donors maintained a normal CD4 + T cell count (Table 2). Interestingly, when we compared length of time on ART, we noted that VQY4910 and MPY1313 were ART suppressed for 206 days and 184 days, respectively, while EXT1011, the cICB non-responsive elite controller, was ART suppressed for 1067 days (Table 2). We also noted that EXT1011 had the lowest CD8+ T cell count and highest CD4:CD8 ratio (Table 2). Whether ART treatment history, CD8+ T cell count, or a combination of these factors explain this donor’s lack of response to cICB is unclear and would require further investigation.

**Table 2.** LVL donors ART suppressed.

|  | CD4+ T count | CD8+ T count | CD4:CD8 ratio | Days on ART |
| --- | --- | --- | --- | --- |
| **VQY4910** | 1025 | 735 | 1.39 | 206 |
| **MPY1313** | 571 | 954 | 0.6 | 184 |
| **EXT1011** | 547 | 341 | 1.6 | 1067 |

## DISCUSSION

CD8+ T cells play a critical role in the elimination of virally infected cells and current efforts aim to fully characterize the molecular mechanisms which regulate their effective immune response to infection, especially in HIV-1 controllers. CD8+ T cell activation upon recognition of an infected cell induces upregulation of inhibitory immune checkpoint receptors (ICRs) which serve to temper the T-cell response during acute infection. During chronic infection, increased upregulation of ICRs results in T-cell exhaustion, a spectrum of progressively impaired effector functions which cause a hierarchical loss of T cell functionality. Our group previously demonstrated that *in vivo* viral control corresponds to *ex vivo* suppression of viral replication in a cohort of HIV-1 controllers (51). In the current study, a cohort of HIV-1 controllers who lost the ability to control viral replication, and were subsequently placed on ART after enrollment, presents a unique opportunity to observe real time changes associated with HIV-1 viral replication and T cell exhaustion within the same subject.

Our data is consistent with the hypothesis that efficient control of viral replication becomes compromised by CD8+ T cell exhaustion. First, we demonstrated that during periods of *in vivo* viral control, ART naïve HIV-1 controllers (LVL) have low expression of CD8+ T cell exhaustion markers and maintain *ex vivo* viral control (Figs 2, 5 and 6). Interestingly, LVLs also demonstrated increased levels of CTLA-4, LAG3 (although these populations were quite rare) and low levels of TIGIT (Fig. 3). This suggests that CD8+ T cells from LVLs may experience an altered form of exhaustion not involving PD-1 and TIM3. Upon *in vivo* loss of control, HIV-1 controllers exhibited upregulated PD-1 and Tim-3 CD8+ T cell exhaustion markers (Fig 4) and lost the ability to suppress *ex vivo* viral replication. This change in exhaustion state persists despite treatment with ART, supporting the idea that ART is not sufficient to reverse T cell exhaustion. We also observed that during periods of control, LVLs exhibited T cell populations similar to NDs, while upon loss of control, T cell populations were similar to HVLs on ART (Fig. 5C, E-G), further suggesting ART does not reverse T cell exhaustion. The incongruous observation that CD8+ T cells express high levels of exhaustion markers despite long-term viral suppression on ART in both LVLs who lost control and HVLs on ART led us to speculate that CD8+ T cell exhaustion in our cohort of HIV-1 controllers mediated the loss of *ex vivo* control.

We evaluated a novel strategy to reverse CD8+ T cell exhaustion through the use of immunotherapy (Fig 8). The loss of viral control in HIV-1 controllers has been attributed to both virological and immunological factors but no mechanism underlying the loss of control has been determined (74–76). It is also important to note that as mechanisms of control themselves are heterogenous, mechanisms of the loss of control are likely also variable. Current literature suggests that both PD-1 and Tim-3 exhaustion markers are associated with decreased CD8+ T cell functionality and their co-expression represents the most severely exhausted phenotype (41, 47, 49, 50). We hypothesized that combinatorial immune checkpoint blockade (cICB) would reverse CD8+ T cell exhaustion in HIV-1 controllers who lost *ex vivo* viral control and restore CD8+ T cell mediated viral control. Consistent with other studies in which blockade of PD-1 or Tim-3 inhibitory immune checkpoint receptors was not sufficient to restore CD8+ T cell effector function alone or resulted in only a modest increase in CD8+ T cell functionality (41, 49, 64, 77, 78), monotherapy was insufficient to restore CD8+ T cell mediated anti-HIV activity in our in vitro assay. However, cICB fully restored CD8+ T cell mediated in vitro viral suppression in CD4+ and CD8+ T cell co-cultures (Figure 8D). This is consistent with other studies which indicate that combinatorial blockade synergistically enhances CD8+ T cell effector functions (64, 77, 78). In contrast, a previous study evaluating the effect of blocking Tim-3 on CD8+ T cells demonstrated that Tim-3+ CD8+ T cells have high cytotoxic potential but are dysfunctional and fail to degranulate (79). They postulated that Tim-3 ICR signaling regulates CD8+ T cell cytotoxic capacities and further demonstrated that in vitro treatment with Tim-3 ICB restored CD8+ T cell anti-HIV activity in ART naïve donors (79). However, this study did not evaluate the co-expression of PD-1 or the loss of control and subsequent ART treatment.

Other studies have evaluated the effects of ICB on T cell isolates but excluded the evaluation of PBMCs. We evaluated the effects of cICB in the context of total PBMCs and observed a varying response in which donors were either cICB responsive or non-responsive (Figure 9). To distinguish between these responses, we compared the exhaustion profiles between these donors (Figure 9). As opposed to the cICB responsive donors, the cICB non-responsive donor had significantly higher expression of PD-1+/Tim-3+ CD8+ T cells, representing the most severely exhausted phenotype. It has been demonstrated that these expression patterns (Tim-3-/PD-1+, Tim-3+/PD-1-, and Tim-3+/PD-1+) are functionally distinct and identify multiple levels of T cell exhaustion (47, 48, 50). This underscores the importance of evaluating the co-expression of both Tim-3 and PD-1 as disease progression can impact the expression of these two ICRs which have been shown to synergistically regulate CD8+ T cell functionality. In addition, our study evaluated ART suppressed donors. It has been demonstrated that treatment with ART alters cytokine signaling which can impact the expression of both Tim-3 and PD-1 on T cells (50). Intriguingly, the cICB non-responsive donor was on ART for a longer time than the cICB responsive donors and was also classified as an elite controller due to an undetectable viral load before beginning therapy (Table 2). This donor also displayed the lowest CD8+ T cell count compared to other donors (Table 2). Although ART can suppress viral replication to undetectable levels, it does not eliminate the virus. It has been demonstrated that viral replication persists despite suppression with ART (68, 80–85). We postulated that despite being ART suppressed, residual viral replication promotes ongoing disease progression, further increasing expression of T cell exhaustion markers which drive the progressive loss of CD8+T cell functionality and development of T cell exhaustion. This study suggests that disease progression has possible implications for differential responses to combinatorial ICB. In addition, our study suggests that Tim-3 and PD-1 have potential for utilization as biomarkers to screen participants with the intent of increasing the efficacy of future clinical trials investigating the use of immunotherapy as a cure strategy for otherwise healthy PLWH.

## MATERIALS AND METHODS

### Sex as a biological variable

Our study examined both male and female human subjects, and similar findings are reported for both sexes. However, the population size presented does not specifically power our study to analyze differences between the sexes and these results may have sex based differences in a larger sample.

### Human subjects

All donors were recruited by the Smith Center for Urban Health and Infectious Disease. Two cohorts of HIV-1 infected donors were recruited, Low Viral Load (LVL) and High Viral Load (HVL) (characteristics at recruitment—Table 1). For the LVL cohort, 5 HIV-1 seropositive ART naïve donors were recruited, who were able to suppress HIV-1 infection independent of known protective HLA alleles. Four of these LVL donors were placed on combination ART during the course of this study. The HVL cohort included HIV-1 seropositive HVL donors pre-and post-therapy. Additionally, we recruited HIV-1 seronegative healthy control donors, Normal Donors that matched the age and characteristics of the two HIV-1 positive groups. For this study, classification as an HIV-1 controller required viremic control for a duration of at least 12 consecutive months in the absence of ART (86–88). HIV-1 controllers are defined as seropositive donors with viral loads of < 2,000 copies/mL and a CD4+ T cell count > 500 cells/mm^3^ (86–88). HIV-1 seropositive donors that had low CD4+ T cell counts, below 200 and had high viral load, above 5,000 copies/mL were designated as HVL donors.

### PBMC isolation and activation

PBMCs were purified from whole blood samples using Ficoll (GE Healthcare) gradient centrifugation and cryopreserved in 90% Fetal Bovine Serum (FBS; HyClone) containing 10% dimethyl sulfoxide (DMSO; Fisher). Frozen PBMCs were thawed and cultured in Roswell Park Memorial Institute (RPMI)-1640 complete media (GenClone) supplemented with 20% heat inactivated FBS, 1x penicillin-streptomycin-glutamine (PSG; ThermoFisher Scientific), and 5% (5 U/mL) human rIL-2 (NIH AIDS Reagent Program). Cells were induced with 5 μg/mL phytohaemagglutinin-P (PHA-P) (Sigma) for 48 h at 37°C and 5% CO_2_.

### CD4+ and CD8+ T-Cell isolation and activation

CD4+ and CD8+ T cells were purified from frozen PBMCs using MACS Miltenyi negative isolation kits (cat# 130-096-533 and cat# 130-096-495, respectively) according to manufacturer’s protocol. Enriched CD4+ and CD8+ T cell populations were independently activated in RPMI complete media with 5μg/mL PHA for 48 h at 37°C and 5% CO_2_.

### HIV-1 virus stock

The HIV-1 stock used in this study was generated by transfecting pNL4-3 (NIH AIDS Reagent Program, ARP-2852, contributed by Dr. M. Martin) into HEK293T cells [American Type Culture Collection (ATCC, CRL-11268)] using TransFectin Lipid Reagent (BioRad, Cat# 1703351) following manufacturer’s instructions. Transfected cells were cultured in Dulbecco’s Modified Eagle’s Medium (DMEM) supplemented with 10% FBS and 1x PSG for 48 h at 37°C and 5% CO_2_. Virus containing supernatants were aspirated from the cells, filtered, and frozen in 1 ml aliquots. Frozen stocks were quantified by p24 Gag ELISA.

### Immune checkpoint blockade spreading infection assay

Isolated activated CD4+T cells were cultured alone as a control. Isolated activated CD4+ and CD8+ T cells were co-cultured 1:1 for 3-6 h at 37°C and 5% CO2. Activated total PBMCs or isolated activated CD4+ and CD8+ T cells were treated with anti-PD-1 (BioLegend, Ultra LEAF purified human anti-PD-1 antibody, clone EH12.2H7) and/or anti-Tim-3 (CD366) antibody (BioLegend, UltraLEAF purified anti-human, clone F38-2E2) immune checkpoint blockade for 24 h at 37°C and 5% CO_2_. Activated treated cells were then infected with 17 ng/ml NL4-3 virus for 24 h at 37°C and 5% CO_2_. After 24h, the virus containing media was removed and replaced. Virus production was evaluated by measuring the p24 levels in the supernatant using ELISA (Zeptometrix). Time points were collected every 48h post infection for 6 to 10 days.

### Flow cytometry

For phenotypic analysis of cell populations, all cells were washed with Phosphate Buffered Saline (PBS) without Ca^2+^ and Mg^2+^ (GenClone) and stained using fluorescently conjugated antibodies against the following cell surface markers in various combinations following manufacturer’s instructions: CD3 (BD Biosciences (BD), Alexa700, clone SP34-2), CD4 (BD, PerCP-Cy5.5), CD8 (BD, BV786, clone RPA-T8), CD279 (PD-1) (Invitrogen, PE-Cyanine7, clone eBioJ105), human TIM-3 (R&D Systems, Alexa Fluor 488, clone 344823), CD223 (LAG-3) (Invitrogen, eFluor 450, clone 3DS223H), TIGIT (Invitrogen, PerCP-eFluor710, clone MBSA43), CD152 (CTLA-4) (Invitrogen, APC, clone 14D3), HLA-DR (BD, APC-H7, clone L243), and CD38 (BD, PE, clone HIT2). Cells were also stained for viability using the live/dead stain Zombie Yellow (BioLegend). Cell enumeration was carried out using Cytek FACSort DxP12 flow cytometer and data analysis using FlowJo v10.6.1 software.

### Multivariate clustering analysis

Clustering and t-Distributed Stochastic Neighbor Embedding (tSNE) dimensionality reduction of flow cytometry data was performed using the FlowSOM and tSNE functions available within FlowJo v10. In brief, the category variable as used to assign an identifier to each individual donor sample to allow examination of specific groups after concatenation and analysis. Cells from each donor sample that were identified as CD3+/Zombie Yellow-singlets were combined into a single data file using the concatenate function with a cutoff of 20,000 events per individual sample. tSNE was performed on this combined data set focusing on TIM3, CD38, TIGIT, PD1, CTLA4, HLA-DR, LAG3, CD8 as variables of interest. Clustering was performed for an assumed 10 clusters (number of stained variables plus two) using FlowSOM and a heatmap was generated. Identified clusters were then mapped back onto the tSNE plot. Selection of the individual donors using the category variable as assigned before concatenization was used to visual groupings by donor type. To examine the expression of individual markers, intensity of staining was mapped onto the tSNE plot as a third dimension.

### Statistics

Data analysis for graphs presented in figures 1, 2, 3, 4, 7 and 8 was performed in GraphPad Prism v10.2.1. Data sets were first analyzed for normality using the Shapiro-Wilk test, then differences were assessed using one-way ANOVA with Tukey’s multiple comparisons test. Data is presented as graphs of all individual data points with a line representing the mean and P-value thresholds as noted for each figure.

### Study approval

The Smith Center for Urban Health and Infectious Disease, East Orange, NJ obtained written informed consent for the collection of blood donations from participating subjects. Samples were collected by trained medical staff under approved University of the Sciences’ protocol (IRB protocol 900702-3 and 797649-3)

### Data availability

Values used to perform analyses and to generate graphs are included in the Supporting Data Values file. Flow cytometry data files will be available through the International Society for the Advancement of Cytometry’s Flow Repository.

## Acknowledgements

We would like to thank Dr. Elias El Haddad for a critical reading of this manuscript and insightful feedback.

## Notes

### Competing Interest Statement

The authors have declared no competing interest.

